# Cross-cultural effects on ensemble coding of emotion in facial crowds

**DOI:** 10.1101/141861

**Authors:** Hee Yeon Im, Sang Chul Chong, Jisoo Sun, Troy G. Steiner, Daniel N. Albohn, Reginald B. Adams, Kestutis Kveraga

## Abstract

In many social situations, we make a snap judgment about crowds of people relying on their overall mood (termed “crowd emotion”). Although reading crowd emotion is critical for interpersonal dynamics, the sociocultural aspects of this process have not been explored. The current study examined how culture modulates the processing of crowd emotion in Korean and American observers. Korean and American participants were briefly presented with two groups of faces that were individually varying in emotional expressions and asked to choose which group between the two they would rather avoid. We found that Korean participants were more accurate than American participants overall, in line with the framework on cultural viewpoints: Holistic versus analytic processing in East Asians versus Westerners. Moreover, we found a speed advantage for other-race crowds in both cultural groups. Finally, we found different hemispheric lateralization patterns: American participants were more accurate for angry crowds presented in the left visual field and happy crowds presented in the right visual field, replicating previous studies, whereas Korean participants did not show an interaction between emotional valence and visual field. This work suggests that culture plays a role in modulating our crowd emotion perception of groups of faces and responses to them.

## Introduction

We routinely encounter groups of people at work, school, or social gatherings. Efficiently reading the emotional states of crowds of people allows us to guide our own reactions and social behaviors. For example, rapidly inferring intent to commit violence from the facial expressions of a mob on the street can allow us to escape in time and avoid potential danger, perhaps by seeking help from another group that looks friendly. Likewise, reading the general mood and receptiveness of an audience allows us to adjust our ongoing behavior, by explaining in more detail or deferring on a point, for a more efficient communication. Such extraction of the prevailing crowd state can occur rapidly and efficiently, by representing the groups of faces as a higher-level description in the form of ensemble coding (Alvarez, 2011; Cohen, Dennett, & Kanwisher, 2016; Haberman & Whitney, 2012). Indeed, recent work on ensemble coding has shown human observers’ remarkable ability to extract average emotion (also termed “crowd emotion”) from sets of faces (e.g., Elias, Dyer, & Sweeny, 2016; Fischer & Whitney, 2011; Haberman, Harp, & Whitney, 2009; Haberman & Whitney, 2007; Hubert-wallander & Boynton, 2015; Im et al., under review; Ji, Chen, & Fu, 2014; Yang et al., 2013), facial identity (de Fockert & Wolfenstein, 2009; Haberman & Whitney, 2007; Leib et al., 2012; Leib et al., 2014; Neumann, Schweinberger, & Burton, 2013), as well as a crowd’s movements (Brunyé, Howe, & Mahoney, 2014; Sweeny, Haroz, & Whitney, 2012) and eye gaze direction (Florey et al., 2016; Sweeny & Whitney, 2014).

Although perceiving average emotion of facial crowds has great social importance in forecasting intentions of groups of people and governing observers’ reactions towards them (e.g., which group would be safe to approach or better to avoid; Im et al., under review), little is known about sociocultural aspects of this process. Only a single study has been done recently to investigate the roles of gender construct in extraction of average identity from a crowd of faces (Bai et al., 2015). However, it still remains unaddressed how different cultural groups (e.g., Westerners and Easterners) perceive crowd emotion when they see faces of same and different races (e.g., Caucasian and East Asian faces). Most of the previous studies on crowd emotion perception have exclusively exploited photographs of Caucasian faces (Fischer & Whitney, 2011; Haberman et al., 2009; Haberman & Whitney, 2007; Hubert-Wallander & Boynton, 2015; Im et al., under review). Although all these studies have been conducted in the Western countries, the specific breakdown of the participants’ ethnic background was not available. We could find only a few exceptions that employed photographs of Asian faces (Chinese faces: Ji et al., 2014; Korean faces: Yang et al., 2013), but only presented to a group of Chinese and Korean participants, respectively. Therefore, none of the existing studies on crowd emotion perception allows us to directly compare how these cultural groups are different in processing crowd emotion from groups of faces of their own and other races.

There is growing interest in empirically testing how differences in cultural experiences shape the way we perceive and respond to the visual world (for a review, Han et al., 2012 and Koelkebeck et al., 2017). Studies comparing Westerners and Easterners have provided convergent evidence on the different perceptual and cognitive styles of those populations (e.g., Lao, Vizioli, & Caldara, 2013; Nisbett & Miyamoto, 2005; Nisbett et al., 2001). For example, Westerners preferably focus on local information, whereas Easterners tend to show perceptual biases towards global information processing in objects (e.g., Masuda & Nisbett, 2001), scene (e.g., Masuda & Nisbett, 2006), and face perception (e.g., Blais et al., 2008; Caldara, Zhou, & Miellet, 2010). Recent evidence also suggests that such cultural differences might rely on different tuning towards visual spatial frequency information across cultural groups (e.g., Miellet et al., 2013). For example, Westerners tend to use preferentially high spatial frequency information from foveal vision, whereas Easterners preferentially process contextual information by relying on extra-foveal vision for face recognition (Miellet et al., 2013), change detection of both low-level visual stimuli (e.g., color blocks; Boduroglu, Shah, & Nisbett, 2009), and scene perception (Masuda & Nisbett, 2001). Such local vs. global contrast between Westerners and Easterners can emerge as early as 80 msec after the onset of a visual stimulus, suggesting that cultural difference is deeply ingrained into the early perceptual processing (Kitayama & Murata, 2013; Lao et al., 2013).

In the current study, we aimed to examine cultural differences in perception of crowd emotion from groups of different faces. We directly compared crowd emotion perception of Westerners (Caucasian and African-American participants) versus Easterners (Korean participants) when they were viewing facial crowds that contained European American faces versus Korean faces. On each trial, we presented two crowds of faces, one in the left and one in the right visual field. We then asked them to choose as quickly and accurately as possible which crowd of faces they would rather avoid. Because participants had to choose one of the two facial crowds presented, their decisions required relative comparison of the two. This allowed us to create task settings that were more representative of the type of social appraisal that we are often involved in on a daily basis: e.g., avoiding a group of strangers who look violent or suspicious in the streets or any other public places.

## Experiment 1

### Methods

#### Participants

A total of 54 undergraduate students in Yonsei University (Seoul, South Korea; N = 25) and in the Pennsylvania State University (State College, PA, USA; N = 29) participated in Experiment 1. All the participants had normal color vision and normal or corrected-to-normal visual acuity. Two participants were excluded from further analyses because they made too many late responses (e.g., RTs longer than 2.5 sec); three participants were excluded because they did not complete the two experimental blocks; one participant was excluded because of too fast responses (e.g., RTs shorter than 0.2 sec) and accuracy lower than chance (50%). Therefore, we ended up with including 48 participants (24 Korean and 24 American). This number of participants we have included in the current study is sufficient providing enough statistical power, given our original study on crowd emotion perception where we have shown the robust and replicable effects from 20-21 participants in each of five successive experiments (Im et al., under review). Thirteen participants were female among 24 Korean participants (11 male) and 12 participants were female among 24 American participants (12 male). Among the 24 American participants, 19 were European American, two were Hispanic, and three were African American. The average age of Korean participants was 20.29 (s.d.: 1.73) years old and the average age of American participants was 18.75 (s.d.: 1.03) years old. The informed written consents were obtained according to the procedures of the Institutional Review Board at Yonsei University and at the Pennsylvania State University for Korean and American participants, respectively. The participants received a course credit for participation in the study.

#### Apparatus and stimuli

Stimuli were generated with MATLAB and Psychophysics Toolbox (Brainard, 1997; Pelli, 1997). In each crowd stimulus, either 4 or 6 morphed faces were randomly positioned in each visual field (right and left) on a white background. Therefore, our facial crowd stimuli comprised either 8 or 12 faces. We used face-morphing software (Norrkross MorphX) to create a set of 51 morphed faces from two highly intense, prototypical facial expressions of the same person for a set of six different identities (3 Korean males and 3 European American males, shown in Figure 1). The Korean face stimuli were taken from the Yonsei Face Database (Yang, Chung, & Chong, 2015), and the American face stimuli were taken from the NimStim Emotional Face Stimuli database (Tottenham et al., 2009). The emotional expression of the faces ranged from 100% happy and 100% angry, with different proportions of morphing between these two (e.g., the neutral with morph of 50% happy and 50% angry faces). The morphed face images were linearly interpolated (in 2% increments) between two extreme faces. Such a morphing approach was adapted from the previous studies on ensemble coding of faces (e.g., Haberman & Whitney, 2007). One of the two crowds in either left or right visual field always had neutral crowd (on average), and the other had emotional crowds on average (e.g., very happy crowd: morphing of angry 32 % and happy 68%) somewhat happy: morphing of angry 40% and happy 60%; somewhat angry: morphing of angry 60% and happy 40%; and very angry: morphing of angry 68% and happy 32%, see Figure 1B). To make an emotional crowd have the mean emotion of one of these morphing ratios, emotionally varying individual faces were randomly chosen from the continuum between 100% angry and 100% happy.

**Figure 1.**
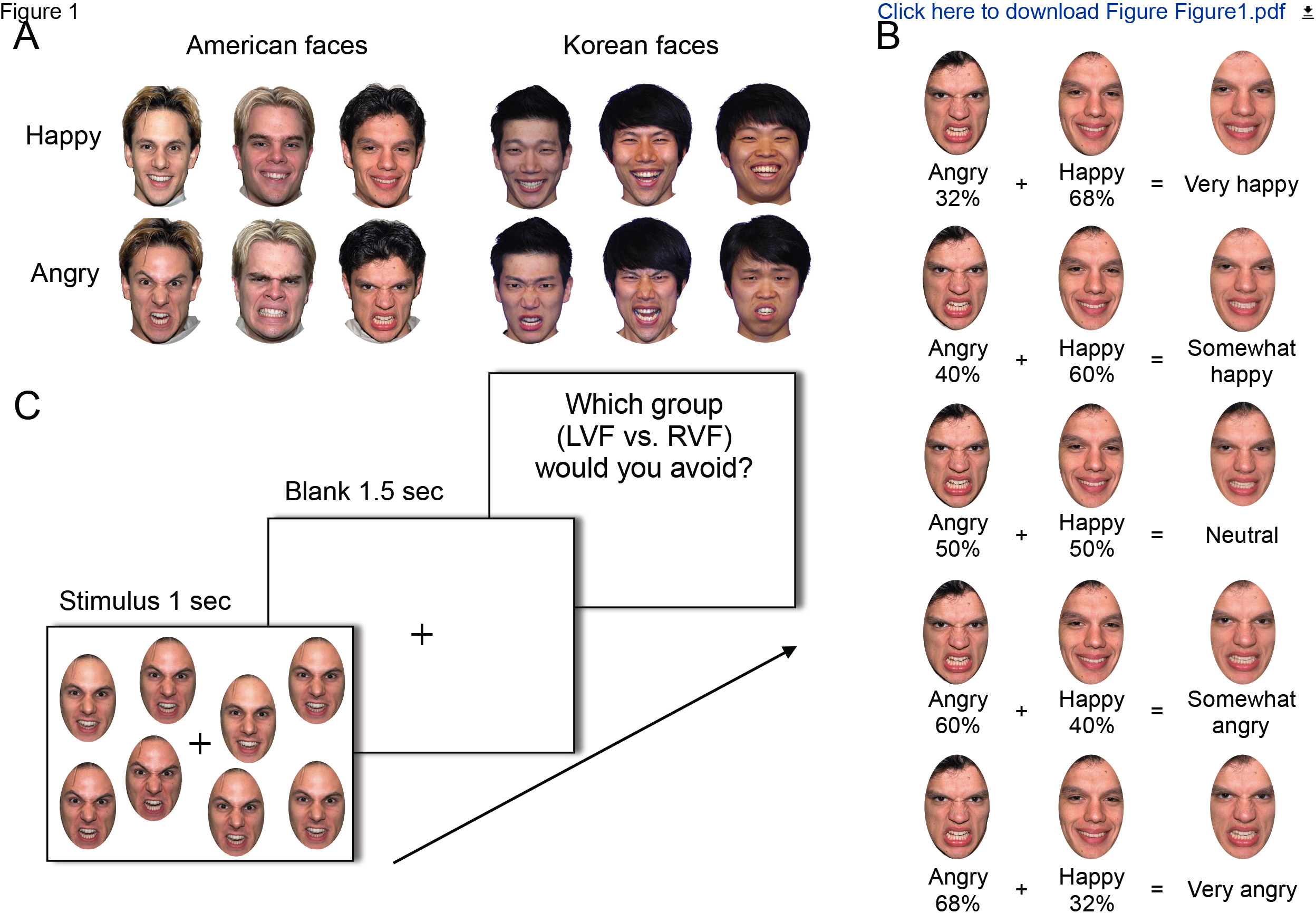
Samples stimuli and a sample experimental trial. (A) Happy and angry faces of three European American faces and Korean faces. (B) Example of different proportions of morphing of the happy and angry faces shown in Figure 1A. Morphing was done for each facial identity. (C) A sample trial.

In order to avoid the possibility that participants simply sampled one or two single faces from each set and compare them to do the crowd emotion task, we ensured that 50% of the individual faces in the neutral set were more expressive than 50% of the individual faces in the emotional sets to be compared. For example, half of the members of the neutral set were angrier than a half of the members of the angry crowd. This manipulation allowed us to assess whether participants used such “sampling strategy” (Myczek & Simons, 2008) rather than extracting the average crowd emotion, because sampling one or two members in a set would only yield 50% of accuracy in this setting. On one half of the trials, the emotional stimulus (i.e., happy or angry) was presented in the left visual field (LVF) and the neutral stimulus was presented in the right visual field (RVF), and it was switched for the other half of the trials. Each face image subtended 2° × 2° of visual angle, and face images were randomly positioned within an invisible frame subtending 13.29° × 18.29°, each in the left and right visual fields. The distance between the proximal edges of the invisible frames in left and right visual fields was 3.70°.

#### Procedure

Figure 1C illustrates a sample trial. We ensured that both Korean and American participants performed the tasks in the same experimental settings and procedures. Participants sat in a chair at individual cubicles and viewed the visual stimuli for 1 second, which was followed by a blank screen for 1.5 second. The participants were instructed to make a key press as soon as possible to indicate which of the two facial crowds on the left or right they would rather avoid. They were explicitly informed that the correct answer was to choose to avoid the facial crowd showing a more negative (e.g., angrier) emotion on average. Responses that were made after 2.5 seconds were considered late and excluded from data analyses. Feedback for correct, incorrect, or late responses was provided after each response. The same written instructions were translated into Korean and English to be provided to Korean and American participants to emphasize that it is critical to make responses as quickly and as accurately as they could.

Korean face stimuli and European American face stimuli were presented in two separate blocks, in a counter-balanced order across participants. Each experimental block had a 4 (different angry/happy morphing ratios: angry 32 % + happy 68%, angry 40% + happy 60%, angry 60% + happy 40%, and angry 68% + happy 32%) × 2 (visual field of presentation, LVF and RVF) x 2 (set size: 8 and 12 faces total) design, with 20 repetitions per condition. This yielded 320 trials total, and the sequence of the trials was randomized within each block.

### Results

Consistent with our previous studies (e.g., Im et al., under review; Yang et al., 2013), we did not find any difference in accuracy (all *p’s* > 0.551) and RT (all *p’s* > 0.773) between different set sizes of the facial crowds (8 vs. 12 faces total; Figures 2A and 2B) in the both cultural groups of participants, suggesting that extraction of crowd emotion does not require serial processing of each individual crowd member, but more likely occurs in parallel. Because there was no effect of crowd size, we collapsed the data from the different crowd size conditions for further analyses.

The overall accuracy for Korean and American facial crowds was 64.57 % (s.d.: 5.13) and 67.17 % (s.d.: 5.83) in Korean participants, and 49.26 % (s.d.: 6.93) and 59.23 % (s.d.: 7.33) in American participants, respectively (Figure 2C). The accuracy of the Korean participants was higher than the chance level (50%) both for Korean facial crowds (*t*(23) = 13.922, *p* < 0.001) and American facial crowds (*t*(23) = 14.443, *p* < 0.001). The performance of the American participants was above chance only for American facial crowds (*t*(23) = 6.170, *t* < 0.001), but not for Korean facial crowds (*t*(23) = 0.5245, *p* = 0.605). Both Korean and American participants showed higher accuracy for American facial crowds compared to Korean facial crowds, and Korean participants were more accurate than American participants both for Korean and American facial crowds. Repeated-measures ANOVA with a within-subject factor of the race of facial crowds and a between-subject factor of the culture of participants (Western vs. Eastern) confirmed these observations with the significant main effects of the race of facial crowds (*F*(1,46) = 27.746, *p* < 0.001) and the culture of the participants (*F*(1,46) = 69.34, *p* < 0.001). The interaction between these factors was also significant (*F*(1,46) = 9.525, *p* < 0.01). Specifically, the interaction showed that American participants were less accurate for Korean facial crowds compared to American facial crowds, whereas Korean participants were equally accurate for both Korean and American facial crowds.

**Figure 2.**
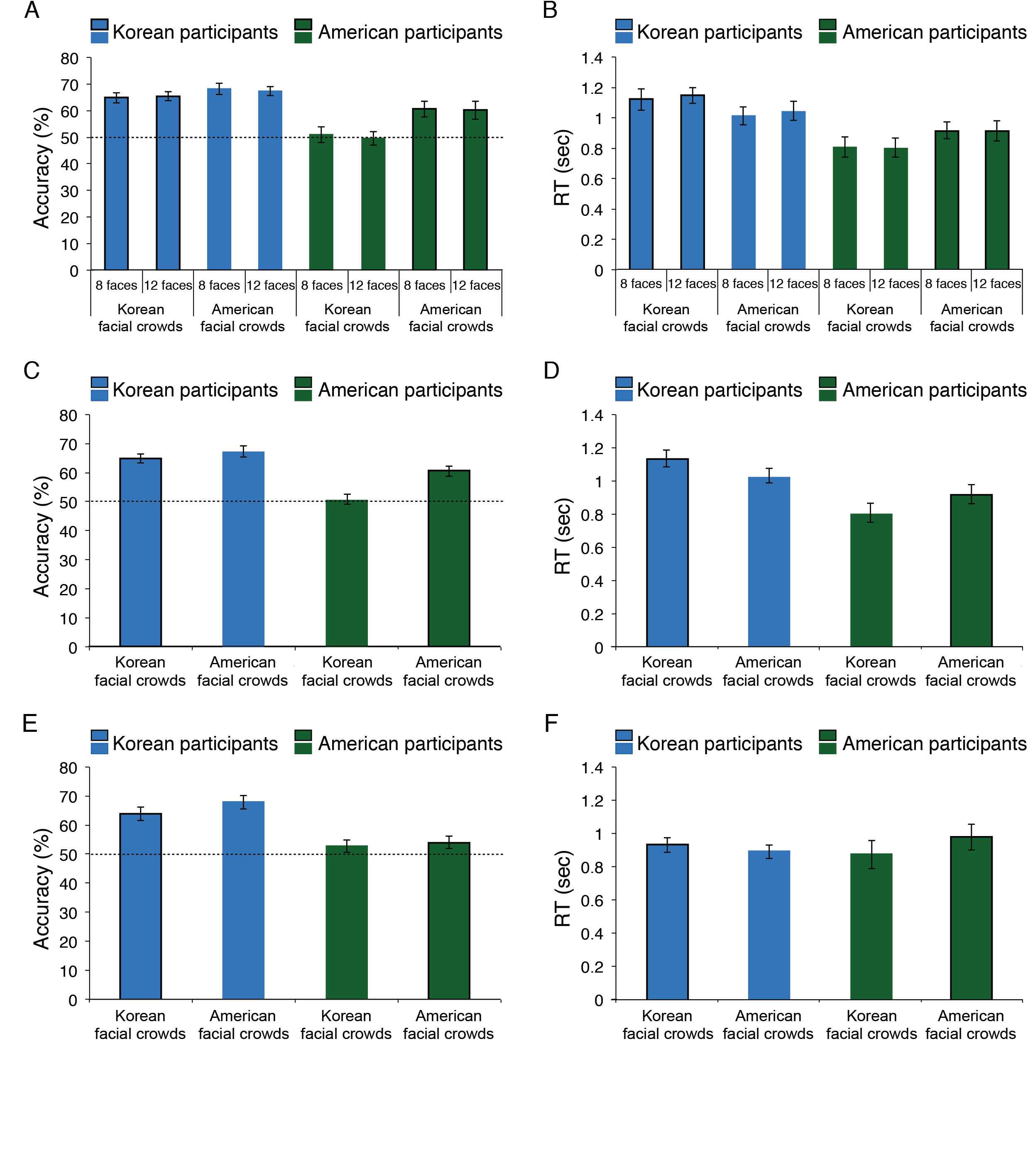
The main results in Experiment 1 (A-D) and in Experiment 2 (E-F). (A) The set size effect on the overall accuracy. The blue bars indicate the accuracy in Korean participants and the green bars indicate the accuracy in American participants. The black outline indicates facial crowds of their own-race for Korean and American participants. The broken like indicates the chance level, 50%. The error bars indicate the standard error of the mean (SEM). (B) The set size effect on the overall RT. (C) Experiment 1: The overall accuracy for Korean and American facial crowds in Korean participants (blue bars) and in American participants (green bars). (D) Experiment 1: The overall RT for Korean and American facial crowds in Korean participants (blue bars) and in American participants (green bars). (E) Experiment 2: The overall accuracy for Korean and American facial crowds in Korean participants (blue bars) and in American participants (green bars). (F) Experiment 2: The overall RT for Korean and American facial crowds in Korean participants (blue bars) and in American participants (green bars).

The median response time (RT) of Korean participants was 1.120 seconds (s.d.: 0.237) for Korean facial crowds and 1.036 second (s.d.: 0.214) for American facial crowds. The median RT for American participants was 0.806 seconds (s.d.: 0.255) for Korean facial crowds and 0.916 seconds (s.d.: 0.273) for American facial crowds (Figure 2D). The main effect of the culture of the participants was significant (*F*(1,46) = 10.95, *p* < 0.01), with Korean participants being much slower than American participants overall. Although the main effect of the race of facial crowds was not significant (*F*(1,46) = 0.232, *p* = 0.632), the interaction between the culture of the participants and the race of facial crowds was statistically significant (*F*(1,84) = 13.143, *p* < 0.001). Specifically, the interaction showed that Korean participants made faster avoidance responses for American facial crowds than for Korean facial crowds whereas American participants made faster avoidance responses for Korean facial crowds than for American facial crowds.

Our recent work has shown the highly replicable effects of hemispheric asymmetry in reading crowd emotion (Im et al., under review): Recognizing the angrier crowd was facilitated when it was presented in the left visual field (LVF) whereas recognizing the happier crowd was facilitated when presented in the right visual field (RVF). Because in the previous study we only recruited American participants and presented them with facial crowds comprising faces from the Ekman face set (Ekman & Friesen, 1976), it is not clear whether this hemispheric asymmetry is universal across different cultures. Therefore, we next examined the patterns of laterality effects in Korean and American participants presented with new Korean and American facial crowd stimuli.

As in the previous study (Im et al., under review), we found an interaction between the visual field and emotional valence of facial crowds in American participants’ responses to American facial crowds. As shown in Figure 3A, the accuracy to detect the angrier crowd (averaged across the two different morphing ratios: angry 60% + happy 40% and angry 68% + happy 32%) was greater when it was presented in the LVF, whereas the accuracy for happier crowd (averaged across the two different morphing ratios: angry 32% + happy 68% and angry 40% + happy 60%) was greater when it was presented in the RVF. Repeated-measures two-way ANOVA with factors of the visual field containing emotional facial crowds (LVF and RVF) and the emotional valence of the crowds (happier and angrier) confirmed a significant interaction of these factors (*F*(1,23) = 14.424, *p* < 0.001). The main effect of emotional valence was also significant (*F*(1,23) = 5.749, *p* < 0.03), but not the main effect of visual field (*F*(1,23) = 0.382, *p* = 0.543). Furthermore, we also found a significant interaction between emotional valence and visual field of presentation with the Korean facial crowd stimuli (*F*(1,23) = 4.919, *p* < 0.04; Figure 3B) which followed the same pattern: greater accuracy for the angrier crowd in the LVF and for the happier crowd in the RVF. Neither the main effect of emotional valence (*F*(1,23) = 0.009, *p* = 0.924) nor the main effect of visual field (*F*(1,23) = 0.006, *p* = 0.980) was significant for the Korean facial crowd stimuli. Therefore, this is not only a direct replication of our previous study in which we showed Ekman faces to American undergraduate participants using a different set of white American faces, but also an extension to a set of other-race facial stimuli (i.e., Korean faces).

**Figure 3.**
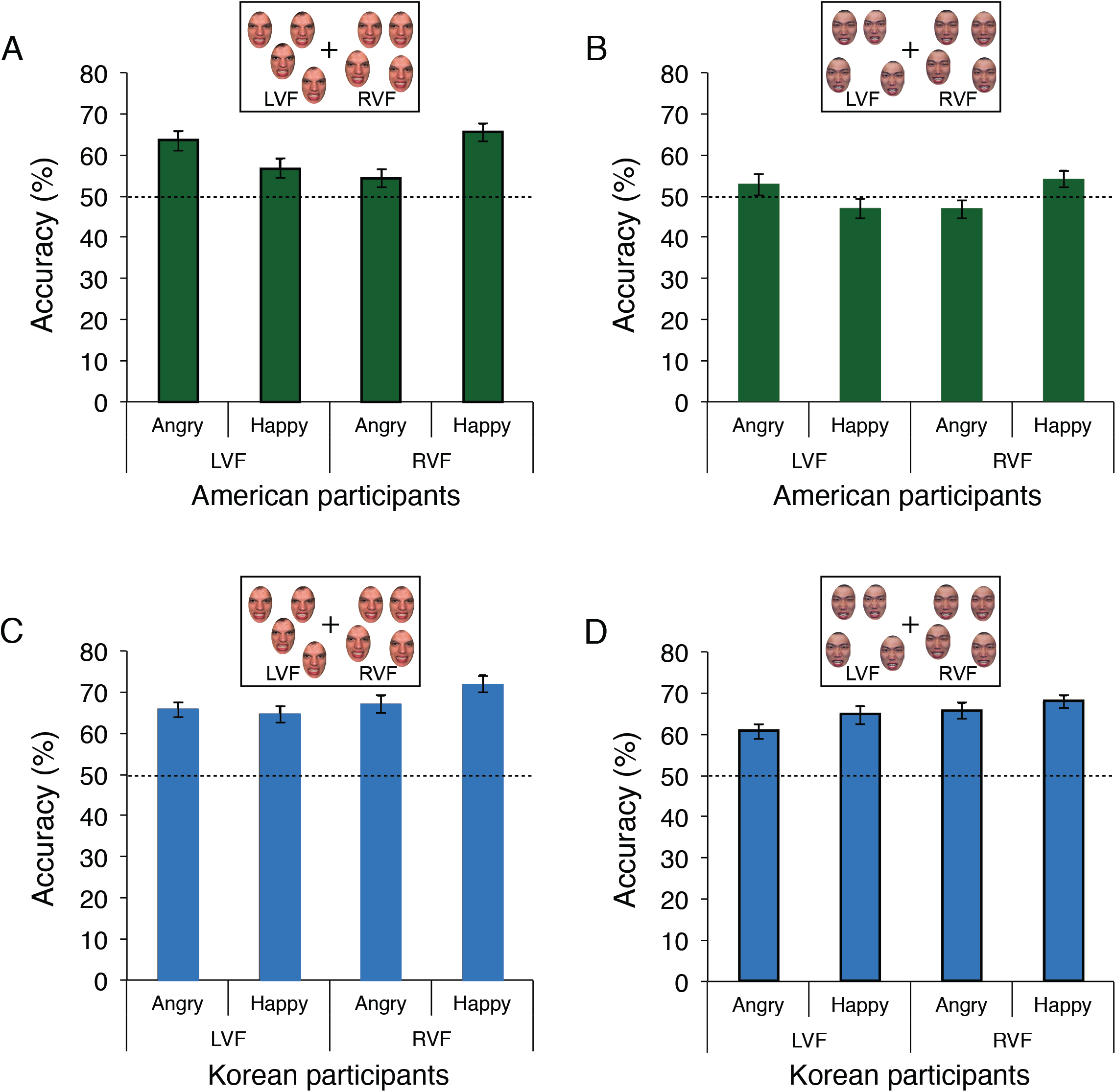
Laterality effect in Experiment 1. (A) The American participants’ accuracy for American facial crowds when they viewed an angrier and a happier facial crowd presented in the LVF and the RVF. The broken like indicates the chance level, 50%. The error bars indicate the standard error of the mean (SEM). (B) The American participants’ accuracy for Korean facial crowds when they viewed an angrier and a happier facial crowd presented in the LVF and the RVF. (C) The Korean participants’ accuracy for American facial crowds when they viewed an angrier and a happier facial crowd presented in the LVF and the RVF. (D) The Korean participants’ accuracy for Korean facial crowds when they viewed an angrier and a happier facial crowd presented in the LVF and the RVF.

We also examined the laterality effect in the Korean participants’ responses to American and Korean facial crowds separately. For American facial crowds (Figure 3C), we observed a similar, but weaker trend in the interaction between emotional valence and visual field, with greater accuracy for angrier crowd in the LVF and happier crowd in the RVF, which was only marginally significant (*F*(1,23) = 3.045, *p* = 0.094). The main effect of visual field was significant (*F*(1,23) = 4.351, *p* < 0.05) with the accuracy for the RVF being greater than for the LVF overall, but the main effect of emotional valence was not significant (*F*(1,23) = 1.195, *p* = 0.286). Unlike for the crowds of European American faces, however, we found that Korean participants did not show the same laterality effect when they viewed Korean facial crowds: The interaction between emotional valence and visual field was not significant (*F*(1,23) = 0.080, *p* = 0.780; Figure 3D). For Korean facial crowds, Korean participants showed greater accuracy for happier crowds than angrier crowds (*F*(1,23) = 8.684, *p* < 0.01) and greater accuracy for the RVF than the LVF (*F*(1,23) = 6.213, *p* < 0.03). Together, American participants seemed to show more pronounced hemispheric lateralization in comparing the emotion of two facial crowds regardless of the race of facial crowds, compared to Korean participants who showed either weak or no laterality effects.

## Experiment 2

In the first experiment, we found differences between Korean vs. American participants in the overall accuracy, RT, and the laterality effects when they made avoidance decisions about emotional crowds of Korean and European American faces. In particular, we observed substantial differences in the overall accuracy and RT, even though we provided both groups of participants with the same instructions (translated into Korean and English) to emphasize both speed and accuracy of responses. Overall, Korean participants made much slower and more accurate responses whereas American participants tended to make much faster but less accurate responses. In Experiment 2, we further tested whether the differences found were due to a speed-accuracy trade-off between the Korean and American participants. In this follow-up study we employed instructions that differentially stressed speed for Korean participants, and accuracy for for American participants, respectively. If the difference in perception accuracy between the Korean and American participants was due to a speed-accuracy trade-off in these groups, stressing speed for Korean participants and stressing accuracy for American participants should reduce the group differences. However, if the differences in perception accuracy and speed we observed in Experiment 1 reflect something else, perhaps different perceptual and cognitive styles, rather than a mere speed-accuracy trade-off, we should still observe differences in accuracy and RT between Korean and American participants.

### Methods

#### Participants

A new group of 46 undergraduate students from Yonsei University (N = 24) and from the Pennsylvania State University (N = 22) participated in Experiment 2. All the participants had normal color vision and normal or corrected-to-normal visual acuity. One participant was excluded from further analyses because of incomplete experimental blocks. Therefore, we ended up including 45 participants (24 Korean and 21 American). Eleven participants were female among 24 Korean participants (13 male) and 10 participants were female among 21 American participants (11 male; 19 European American and 2 African American). The average age of Korean participants was 20.08 (s.d.: 2.30) years old and the average age of American participants was 18.86 (s.d.: 1.06) years old.

#### Apparatus and stimuli

All the aspects of the apparatus and the stimuli were identical to those in Experiment 1.

#### Procedure

All the aspects of the experimental procedure were identical to those in Experiment 1, except that the written instructions provided to Korean and American participants were revised. For Korean participants, speed was particularly emphasized over accuracy whereas for American participants, accuracy was particularly emphasized over speed. The actual instructions given to Korean and American participants are shown as follow:

*<u>a. Instruction for Korean participants (translated into Korean in the actual experimental session):</u> On each trial, two crowds of faces with varying emotional expressions will be presented for 1 second. Your task is to choose which of the two groups of people (on the left or right) you would rather avoid. Press ‘F’ to avoid left group, and press ‘J’ to avoid right group on the screen. Correct answer is to choose angrier (more negative) crowd on average. Feedback will be provided on each trial. Please try to remain as accurate as possible. However, we require fast responses for our analyses: Accuracy at the expense of speed will undermine our study, because responses that are too slow cannot be analyzed. Please rely on your first impression of the stimuli and try to make response as quickly as you can*.
*<u>b. Instruction for American participants:</u> On each trial, two crowds of faces with varying emotional expressions will be presented for 1 second. Your task is to choose which of the two groups of people (on the left or right) you would rather avoid. Press ‘F’ to avoid left group, and press ‘J’ to avoid right group on the screen. Correct answer is to choose angrier (more negative) crowd on average. Feedback will be provided on each trial. Please try to remain as fast as possible. However, we require accurate responses for our analyses: Speed at the expense of accuracy will undermine our study, because only accurate responses can be analyzed. Please try to make response as accurately as you can.*

## Results

### Korean participants: stressing response speed

The median RTs in Experiment 2 were 0.944 second (s.d.: 0.178) for Korean facial crowds and 0.911 second (s.d.: 0.171) for American facial crowds (Figure 4A). In order to test whether the different instruction stressing the response speed was effective, we compared the RTs obtained in Experiment 2 with those in Experiment 1. Repeated measures ANOVA with a within-subjects factor of the race of facial crowds (European American faces vs. Korean faces) and a between-subjects factor (Experiment 1 vs. Experiment 2) revealed significantly faster RTs for Experiment 2 than Experiment 1 (*F*(1,46) = 7.340, *p* < 0.01) and significantly faster RTs for American facial crowds than Korean facial crowds (*F*(1,46) = 11.332, *p* < 0.01, although the interaction was not significant (*F*(1,46) = 2.161, *p* = 0.148). Thus, our new instruction that emphasized the response speed was effective resulting in faster responses of Korean participants in Experiment 2.

**Figure 4.**
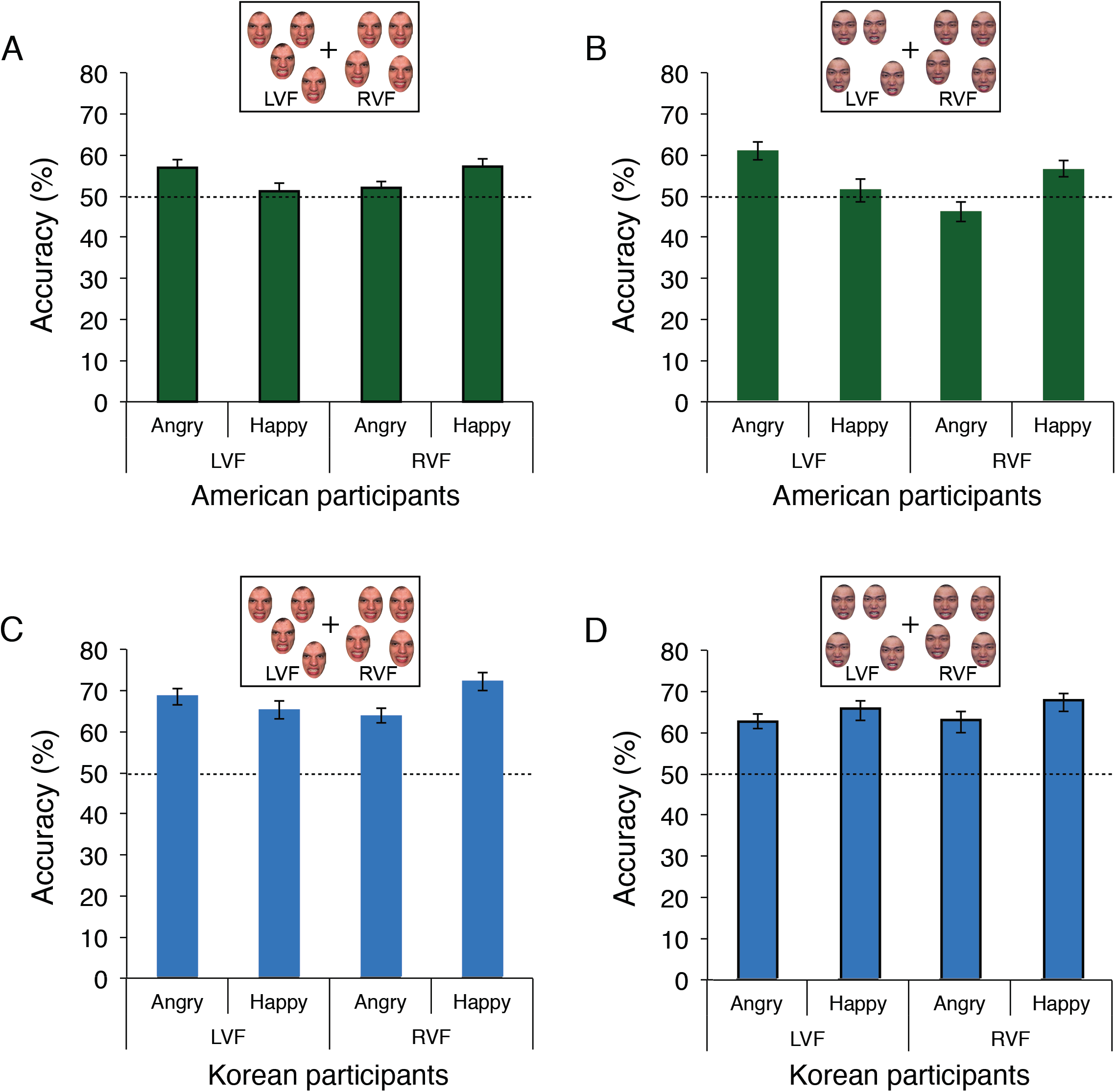
Laterality effect in Experiment 2. (A) The American participants’ accuracy for American facial crowds when they viewed an angrier and a happier facial crowd presented in the LVF and the RVF. The broken like indicates the chance level, 50%. The error bars indicate the standard error of the mean (SEM). (B) The American participants’ accuracy for Korean facial crowds when they viewed an angrier and a happier facial crowd presented in the LVF and the RVF. (C) The Korean participants’ accuracy for American facial crowds when they viewed an angrier and a happier facial crowd presented in the LVF and the RVF. (D) The Korean participants’ accuracy for Korean facial crowds when they viewed an angrier and a happier facial crowd presented in the LVF and the RVF.

Despite **the faster RT’s for** Korean participants in Experiment 2, however, their accuracy did not decrease significantly (Figure 4B). Repeated-measures ANOVA showed that **Korean participants’** accuracies in Experiments 1 and 2 were not significantly different (*F*(1,46) = 0.825, *p* = 0.329). Therefore, stressing response speed did not seem to impair **Korean participants’ response accuracy**. We also found that the accuracy for American facial crowds was significantly greater than for Korean facial crowds overall (*F*(1,46) = 29.258, *p* < 0.001), replicating the results of Experiment 1.

### American participants: stressing response accuracy

In Experiment 1, we observed that American participants yielded much faster, but less accurate, responses compared to Korean participants. Thus, we asked a new group of American participants to try to spend enough time to view the stimuli and respond as accurately as possible in Experiment 2. However, the new instructions emphasizing accuracy over speed did not seem to improve American participants’ accuracy. The accuracy was 52.71% (s.d.: 6.777) for Korean facial crowds and 53.44% (s.d.: 7.319) for American facial crowds (shown in Figure 2E). Repeated-measures ANOVA revealed that the accuracy in Experiment 2 was not significantly different from that in Experiment 1 (*F*(1,43) = 1.917, *p* = 0.173), although the main effect of the race of facial crowds (*F*(1,32) = 14.02, *p* < 0.001) and the interaction (*F*(1,32) = 10.46, *p* < 0.01) were significant. We found a similar pattern for RT’s of American participants in Experiment 1 vs. Experiment 2: The RT’s for Experiment 1 were not significantly different from those for Experiment 2 (*F*(1,43) = 1.285, *p* = 0.263); and the RTs for Korean facial crowds were faster than for American facial crowds (Figure 2F). Thus, we conclude that the instructions to American participants to focus on accuracy instead of speed failed to shift the participants’ speed-accuracy trade-off criterion and did not significantly influence American participants’ accuracy or RT.

Finally, we tested the laterality effects in making avoidance decision between two facial crowds when different instructions stressing either speed or accuracy were given to participants. As shown in Figures 4A and 4B, we found the same laterality effect when American participants viewed American facial crowds and Korean facial crowds that we observed in Experiment 1. Both for American and Korean facial crowds, American participants in Experiment 2 again were more accurate to detect angrier crowds in the LVF, and happier crowds in the RVF, confirmed by a repeated-measures ANOVA with significant interaction between two factors of the visual field and the emotional valence (European American faces: F(1, 20) = 4.692, *p* < 0.05; Korean faces: *F*(1,20) = 4.370, *p* < 0.05). Unlike American participants, Korean participants showed different patterns of laterality effects for European American facial crowds and for Korean facial crowds: For European American facial crowds, Korean participants were also more accurate to detect angrier crowds presented in the LVF and happier crowds presented in the RVF (the interaction was marginally significant: *F*(1,23) = 3.250, *p* = 0.08; Figure 4C). However, Korean participants did not show significant interaction between visual field and emotional valence (*F*(1,23) = 0.008, *p* = 0.931) or main effects of visual field (*F*(1,23) = 0.053, *p* = 0.819) and emotional valence (*F*(1,23) = 1.456, *p* = 0.240; Figure 4D).

## Discussion

The current study examined cultural differences between American and Korean participants in processing crowd emotion of faces belonging to either their own or other racial group. Here we report three main findings: 1) Korean participants were more accurate overall than American participants in choosing the angrier facial crowd to avoid, for both Korean and European American faces; 2) Korean participants made faster avoidance responses for American facial crowds than for Korean facial crowds, whereas American participants made faster responses to Korean facial crowds than to white American facial crowds; 3) American participants showed highly lateralized response patterns in which they were more accurate to detect angrier facial crowds presented in the LVF, and happier facial crowds presented in the RVF, whereas Korean participants did not show a robust pattern of hemispheric lateralization, particularly for own-race faces.

The better overall accuracy in Korean participants is not simply due to a speed-accuracy trade-off. Although Korean participants were much slower than American participants in Experiment 1, the instructions to Korean participants to focus on speed in Experiment 2 lowered their RTs to be on par with those of American participants, while maintaining the same higher level of accuracy. Conversely, American participants were much less accurate than Korean participants both in Experiments 1 and 2, even when they were instructed to focus on making correct responses. Instead, we suggest that the better accuracy in Korean participants than American participants for perceiving crowd emotion of groups of faces may be attributable to better sensitivity of Easterners to global information extracted by holistic processing (Abel & Hsu, 1949; Ji, Peng, & Nisbett, 2000; Kitayama et al., 2003; Masuda & Nisbett, 2001; Nisbett, 2003; Nisbett & Miyamoto, 2005; Norenzayan & Nisbett, 2000; Peng & Nisbett, 1999). Such tendency can generalize to social contexts, suggesting that Westerners tend to see emotions as individuals’ feelings seen collectively, whereas Easterners tend to see them as a group feeling. For example, Masuda et al. (2008) presented a set of cartoon pictures of an individual surrounded by a group of four other people and asked Japanese and Western participants to judge the emotion of the central person. They found that the emotion of surrounding people influenced judgment on the emotion of the central person in Japanese, but not in Western participants, and Japanese participants looked at the people surrounding the central person more than Westerners did. Moreover, our finding is also in line with the previous studies that showed American participants perform better in the absolute task whereas Japanese perform better in the relative task (Hedden et al., 2008; Kitayama et al., 2003) and that Chinese participants are better at detecting cooccurrence and co-variation of events compared to American participants (Ji et al., 2000).

We found that both American and Korean participants made faster avoidance responses when they viewed facial crowds of other races (e.g., American participants being faster for Korean facial crowds and Korean participants being faster for American facial crowds). Such a speed advantage for other-race faces has been previously observed in single face classification and recognition of human subjects (Caldara et al., 2004) and similar attentional bias toward conspecifics’ emotions in primates (Kret et al., 2016). For example, by presenting Caucasian participants with single faces of Caucasians versus East Asians, Caldara et al., (2004) found that other-race faces (e.g., Asian faces) are classified faster than same-race faces (e.g., Caucasian faces). They suggested that typically lesser experience with other-race faces engenders fewer semantic representations, which in turn increases the speed of processing for such stimuli. Moreover, it has been suggested that people tend to be more favorable, in terms of affective reactions and resource allocation, to members of their own group (in-group) than to other groups (Brewer, 1979; Tajfel & Turner, 1986). Allocating fewer resources to detailed features in out-group faces would also lead to insufficient individuation of members of the group, as it is known as the “out-group homogeneity effect” (Linville, Salovey, & Fischer, 1986; Messick & Mackie, 1989; Mullen & Hu, 1989; Park, Judd, & Ryan, 1991; Quattrone, 1986; Wilder, 1986). These factors could have contributed to our new findings of faster avoidance responses to facial crowds of other-race faces in both American and Korean participants. Therefore, our findings provide a conceptual replication and extension of theories of group perception, specifically defined by race, in more naturalistic social settings where observers perceive groups of emotional faces and make an affective evaluation between the groups.

Hemispheric asymmetry allows for processing of incoming perceptual inputs in parallel for competing goals (Rogers, Vallortigara, & Andrew, 2013). Goal-dependent parallel processing is particularly useful when a large number of complex stimuli (such as a crowd of emotional faces) and competing cognitive goals tax the processing capacity of the visual system, as was the case in our task. For example, the right hemisphere (RH) is suggested to dominate in attending to novelty, detecting behaviorally goal-relevant, clear sensory events, executing rapid responses, and extracting global features, whereas the left hemisphere (LH) is suggested to control responses for consideration of alternatives, resolve ambiguities, and process local features by inhibiting RH (Corbetta & Shulman, 2002; Rogers, 2002; Robertson, Lamb, & Knight, 1988). Recently, we have shown that the pattern of hemispheric lateralization in reading crowd emotion is dependent on the ongoing task goal in the context of social decision (Im et al., under review). When participants made avoidance decisions choosing between two facial crowds, they were more accurate for angrier crowds presented in the left visual field (LVF) and happier crowds presented in the right visual field (RVF). When they made approach decisions, however, the pattern was reversed: They were more accurate for happier crowds presented in the LVF and angrier crowds presented in the RVF.

Because Im et al. (under review) presented American participants only with Caucasian faces selected from the Ekman face set (Ekman & Friesen, 1976), it was unclear if this effect could be generalized to other cultural groups. The current study not only replicates their findings by using different stimuli (both for Caucasian and Korean facial crowds) in a different cohort of American participants, but also shows that Korean participants exhibit different patterns of hemispheric lateralization from American participants.

The differences in hemispheric lateralization between American participants (e.g., highly lateralized) and Korean participants (e.g., weakly or not lateralized) may reflect the different breadth of attention that results in different styles of engaging in the task and achieving the task goals. Westerners are suggested to be more narrowly focused, whereas Easterners have been suggested to have a broader focus or be more holistic in the application of attention to perceptual objects (Nisbett et al., 2001). Kitayama & Murata (2013) suggested that focused attention, more dominant in Westerners than Easterners might be due to their strong independent orientation and corresponding tendency to pursue an action towards a goal-relevant object from the very beginning of processing (Kitayama & Murata, 2013; Markus & Kitayama, 1991; Varnum et al., 2010). Conversely, global attention for holistic processing, dominant in Easterners, might be due to their tendency to be receptive to a wider range of incoming pieces of information and adjust their behaviors accordingly (Kitayama & Murata, 2013). Given the LVF advantage for detecting clear and goal-relevant features (e.g., angry crowds in our avoidance task) and RVF advantage for analysis of the alterative and ambiguous features (e.g., happy crowd in our avoidance task), the pattern of lateralization that we observed in American participants may reflect their tendency towards goal-oriented, focused processing to maximize the efficiency by selectively facilitating processing of angrier crowds in the LVF and happier crowds in the RVF, but instead compensating processing of other combinations (happier crowds in the LVF and angrier crowds in the RVF). Our findings provide novel empirical evidence for the cross-cultural variations in hemispheric lateralization that is closely associated with socio-affective decision making in different cultural groups.

Previous studies have also reported qualitatively different patterns of hemispheric lateralization between Westerners and Easterners. To illustrate, Western subjects showed LVF superiority while Japanese subjects showed visual field symmetry in response to geometric shapes (Hatta & Dimond, 1980); Westerners showed bilateral activity to face stimuli in the fusiform face area (FFA) whereas Easterners showed more right lateralization (Goh et al., 2010); and Westerners showed left hemisphere (LH) dominance whereas Easterners showed right hemisphere (RH) dominance in the baseline activation (Moss, Davidson, & Saron, 1985) during the EEG resting state (with eyes-closed). Although the results are mixed, allowing for only limited conclusions, these findings along with our own findings at least suggest that there may be cross-cultural differences in hemispheric lateralization that reflect ethnic characteristics in behaviors, such as biases in perception, attention, cognition, and social attribution.

### Limitations of the current study

Before concluding, it is worth noting that our findings should be interpreted with caution. Although our findings are consistent with the well-accepted framework on two distinct but culturally dependent cognitive styles of information processing (analytical vs. holistic), different explanations are also possible, due to a variety of practical constraints and confounding factors associated with cross-cultural experimental research. We wish to acknowledge three possible difficulties in interpretation.

First, it may be the case that Korean and American participants have used different implicit rules and criteria regarding the balance between speed and accuracy. Although we manipulated the test instructions and showed that the differences in accuracy and RT between Korean and American participants did not merely reflect differences in the speed-accuracy trade-off, we cannot completely rule out Korean and American participants’ different tendency to emphasize one aspect over the other regardless of the instruction. American norms for timed measures typically remain within normal limits for a single error, suggesting a relatively specific speed-accuracy trade-off valued in American culture (Strutt et al., 2015). However, some other cultural groups suggest that speed and quality are contradictory and highlight the relative importance of accuracy over speed, for a good product is the result of a slow and careful process (Ardila, 2007). Our results may partly support this possibility in that Korean participants did not sacrifice their accuracy for speed when they were instructed to focus on making fast responses while American participants did not slow down their responses when they were instructed to spend more time to be more accurate. Future work should empirically address the cross-cultural differences in the criteria for speed-accuracy trade-off for better understanding of the cultural influences on various perceptual and cognitive tasks.

Second, it is possible that the better overall accuracy in Korean participants than in American participants results from the type of the task we employed: avoiding one of the two emotional facial crowds. It has been shown that North Americans tend to be more attentive to approach-oriented information, whereas East Asians tend to be more attentive to avoidance-oriented information. For example, Hamamura et al. (2009) showed that American participants recalled more items describing approach-motivating events (e.g., gorgeous weather for hiking) whereas Japanese recalled more items describing avoidance-motivating events (e.g., stuck in a traffic jam). Moreover, it has been shown that Asian Americans and Koreans were more likely to embrace avoidance personal goals relative to European Americans (Elliot et al., 2001); Americans rated a tennis game that was framed as an opportunity to win as more important than one that was framed as an opportunity to avoid a loss, whereas Chinese participants showed the reverse pattern (Lee, Aaker, & Gardner, 2000); and Canadian participants were motivated by success feedback more than failure feedback whereas Japanese participants were motivated by failure feedback than success feedback (Heine, Ide, & Leung, 2001; Oishi & Diener, 2003). Likewise, the better accuracy for Korean participants than Americans in the current study may be also associated with the avoidance task that we employed.

In conclusion, the current study reports cross-cultural differences between American and Korean participants in reading crowds emotion of facial groups. Cultural considerations in group perception are a rarely investigated topic although it seems that differences between countries in social norms may be closely related to their interpersonal perception and interactions. Our work offers an initial inquiry into cross-cultural differences in ensemble perception of faces with great social significance by presenting the same visual stimuli to two different cultural groups. Our work is a small step to bridge a gap between literature on ensemble coding of facial stimuli and a burgeoning area of cross-cultural work showing the importance of culture in understanding social perceptual processes.

## Acknowledgments

This work was supported by the National Institutes of Health R01MH101194 to K.K. and to R.B.A., Jr., and by the National Research Foundation of Korea (NRF) grant funded by the Korea government (NRF-2016R1A2B4016171) to S.C.C.

Kestas Kveraga: Reginald B. Adams, Jr: Sang Chul Chong

Data collection was conducted at the Pennsylvania State University for American participants and at Yonsei University for Korean participants. Informed written consent was obtained in all studies according to the procedures of the Institutional Review Board at the Pennsylvania State University and at Yonsei University. The participants received a course credit for their participation.

Troy G. Steiner: Jisoo Sun

## Author Contributions

H. Y. Im, R. B. Adams, and K. Kveraga developed the study concept and all the authors contributed to study design. Testing and data collection were performed by J. Sun, T. G. Steiner, and D. N. Albohn. H. Y. Im analyzed the data and all the authors wrote the manuscript.

